# Individual Differences in Speech Monitoring: Functional and Structural Correlates of Delayed Auditory Feedback

**DOI:** 10.1101/2025.10.22.683659

**Authors:** Muge Ozker, Laura Giglio, Ahmad Beyh, Stephanie J. Forkel, Peter Hagoort

**Author notes:** Equal contribution (Stephanie J. Forkel, Peter Hagoort).

## Abstract

Sensory feedback is essential for the fine-tuning of motor actions, and speech production is no exception. It depends on continuous self-monitoring to ensure that produced sounds match intended targets. Delaying auditory feedback (DAF) disrupts this alignment and impairs fluency, providing a powerful tool to investigate sensorimotor control. We combined functional and diffusion-weighted MRI in 31 participants performing a word-production task under delayed (DAF) and immediate (no-DAF) auditory feedback. While all participants slowed their speech under DAF, the extent of this effect varied across individuals and was quantified using a susceptibility index (SI). At the group level, DAF elicited increased activation in a right-lateralized network encompassing the superior temporal gyrus, supramarginal gyrus, inferior frontal gyrus, supplementary motor area, and left cerebellum. Incorporating individual differences revealed that higher susceptibility was associated with greater activation in left-hemisphere speech motor homologues and larger volume of the right long arcuate fasciculus, a white-matter pathway connecting auditory and motor speech regions. This pattern suggests that vulnerability reflects increased recruitment of neural resources and a stronger reliance on auditory-motor coupling. In contrast, resilience was associated with greater engagement of the bilateral angular gyrus and higher fiber density in the right posterior arcuate fasciculus, which connects auditory and somatosensory speech regions. This finding indicates that resilience is supported by a posterior circuit that efficiently integrates multi-modal sensory feedback. Together, these findings link functional dynamics with underlying structural connectivity to reveal how a right-lateralized network supports speech control, while accounting for individual differences in susceptibility to fluency disruption.

**Significance Statement:** Fluent speech depends on the brain’s ability to monitor self-produced sounds and sensations from articulatory organs to adjust motor commands in real time. To uncover the neural basis of this process, we combined a fluency-disrupting paradigm, delayed auditory feedback (DAF), with functional and structural neuroimaging. This multimodal approach revealed that while DAF processing relies on a right-lateralized network, more susceptible individuals show enhanced recruitment of left-hemisphere monitoring regions. We also found that stronger white-matter connections in the right posterior arcuate fasciculus predict greater resilience and fluency. These findings provide an anatomically grounded account of how auditory and somatosensory feedback interact to support speech production, offering new insight into why some individuals are more susceptible to fluency breakdowns and related disorders.

## INTRODUCTION

During speech production, the brain continuously monitors how closely the sounds being produced match the intended outcome (1). Hearing one’s own voice is central to this monitoring process, as auditory feedback enables the detection and correction of speech errors in real time (2, 3), ensuring that motor commands produce the desired speech sounds. Disrupting auditory feedback can alter speech output, as many people have experienced when speaking while listening to loud music through headphones, often raising their voice unintentionally because they cannot accurately regulate its loudness (4, 5).

To investigate the neural mechanisms underlying this monitoring system, researchers often manipulate auditory feedback during speech production to introduce controlled mismatches between expected and perceived sounds. Such perturbations simulate speech errors and reveal how the brain adjusts ongoing production to maintain accuracy and fluency. Depending on the nature of the manipulation, different aspects of speech are disrupted, and distinct corrective strategies are engaged. Spectral perturbations, such as shifts in fundamental frequency (F0) or formant frequencies (F1 and F2), typically elicit directional compensatory adjustments that serve to minimize the perceived acoustic error. For example, shifting voice pitch often triggers a compensatory change in the opposite direction (2, 6, 7), while formant manipulations prompt speakers to reconfigure their articulatory posture to reach the intended vowel target (8–12). In contrast, when the timing of auditory feedback is altered, as in delayed auditory feedback (DAF), the monitoring system must resolve a temporal asynchrony between articulation and perception. Under these conditions, speakers commonly slow their speech rate, prolong vowels, or reset syllables to realign their production with the delayed auditory signal (13–15). Taken together, these varied adjustments illustrate the brain’s ability to resolve sensorimotor discrepancies by either correcting specific acoustic features or restructuring the temporal flow of speech to maintain coordination.

Most neuroimaging studies on speech monitoring have used pitch (16–22) or formant perturbations (3, 23, 24), which mainly probe brain mechanisms underlying vocalization and articulation control. In contrast, DAF processing remains comparatively underexplored, leaving temporal coordination processes that are fundamental to fluent speech unclear. The limited PET and fMRI literature on DAF has employed a range of methodological approaches, including various delay durations (50, 100, or 200 ms), different baseline conditions (e.g., contrasting DAF with rest or with immediate auditory feedback), trial structures (randomized versus blocked), and task demands (e.g., speaking at different rates). Despite these methodological differences, early investigations consistently reported the strongest disruption of speech at 200 ms delay and increased activation in the bilateral superior temporal gyrus (STG) in response to delayed feedback, while showing little or no reliable engagement of motor-related regions (25–27). More recently, however, an fMRI study from Agnew et al. (28) and electrocorticography (ECoG) work from Ozker et al. (29), both directly contrasting randomized delayed and immediate feedback conditions, have demonstrated that DAF processing recruits a substantially broader network encompassing inferior parietal and frontal cortices in both hemispheres. These later findings indicate that DAF engages more than auditory error detection alone. However, a comprehensive whole-brain account of the neural substrates supporting the transition from auditory error detection to motor adjustment, including functional lateralization and the role of deeper structures, has yet to be fully characterized.

In addition, individuals vary widely in their susceptibility to altered auditory feedback, yet the neural substrates underlying this variability also remain poorly understood. Clinical studies highlight striking differences: DAF induces more dysfluencies in individuals with schizophrenia (31) or autism spectrum disorder (32) than in neurotypical controls, whereas patients with conduction aphasia show reduced susceptibility (33). Intriguingly, while DAF often elicits stutter-like speech in neurotypical speakers, it can improve fluency in people who stutter (34). Even within neurotypical populations, substantial interindividual variability in DAF susceptibility has been reported (28, 35–37), yet its neural basis remains unknown.

Current models of speech production describe speech monitoring as a dynamic interaction between perisylvian motor and sensory regions through feedforward and feedback processes (38, 39). However, our understanding of this process is shaped largely by cortical activation patterns observed in neuroimaging studies, with comparatively little knowledge on the contribution of subcortical structures and the white matter pathways that integrate the different components of the speech network. A central white matter pathway that arches around the lateral fissure, and link the frontal motor, parietal somatosensory, and temporal auditory regions relevant for speech production is the arcuate fasciculus (AF). The AF has long been recognized as critical for healthy speech and language processing (40–42) and implicated in language disorders like aphasia (43–45) and auditory hallucinations in schizophrenia (46). Anatomically, the AF can be subdivided into three components: a direct long fronto-temporal segment, and two indirect fronto-parietal and parieto-temporal tracts that link these regions via the inferior parietal lobe (41). Different segments of the AF have been linked to distinct components of speech and language processing, including fluency (47), naming (40), repetition (45), and lexical-semantic processing (48). However, their specific contributions to sensory feedback control of speech remain unclear. Notably, the AF shows marked interindividual variability in its anatomical structure (e.g. presence/robustness of the long segment, lateralization, fiber capacity) (49, 50), raising the question of whether this structural variability contributes to individual differences in susceptibility to DAF.

To address these questions, we conducted a speech production experiment in neurotypical individuals using DAF during *f*MRI, alongside diffusion-weighted imaging (DWI) and in-scanner voice recordings. Our objectives were to: (1) identify cortical networks involved in monitoring speech coordination and fluency through DAF manipulation, (2) determine the role of the AF in supporting this process, and (3) establish the neural substrates that underlie individual differences in susceptibility to DAF measured by the degree of speech slowing.

## RESULTS

### Behavioural Results

For the behavioral analysis, we examined participants’ voice recordings acquired during scanning by calculating word duration for each trial. Speaking with 200 ms delayed auditory feedback (DAF) slowed speech rate compared to speaking with immediate feedback (no-DAF), as reflected by a significant increase in word duration across participants (no-DAF: 0.85 ± 0.02 s; DAF: 0.95 ± 0.03 s; paired t-test: t = 7.859, p < 0.001; **Fig. 1A**).

**Fig. 1.**
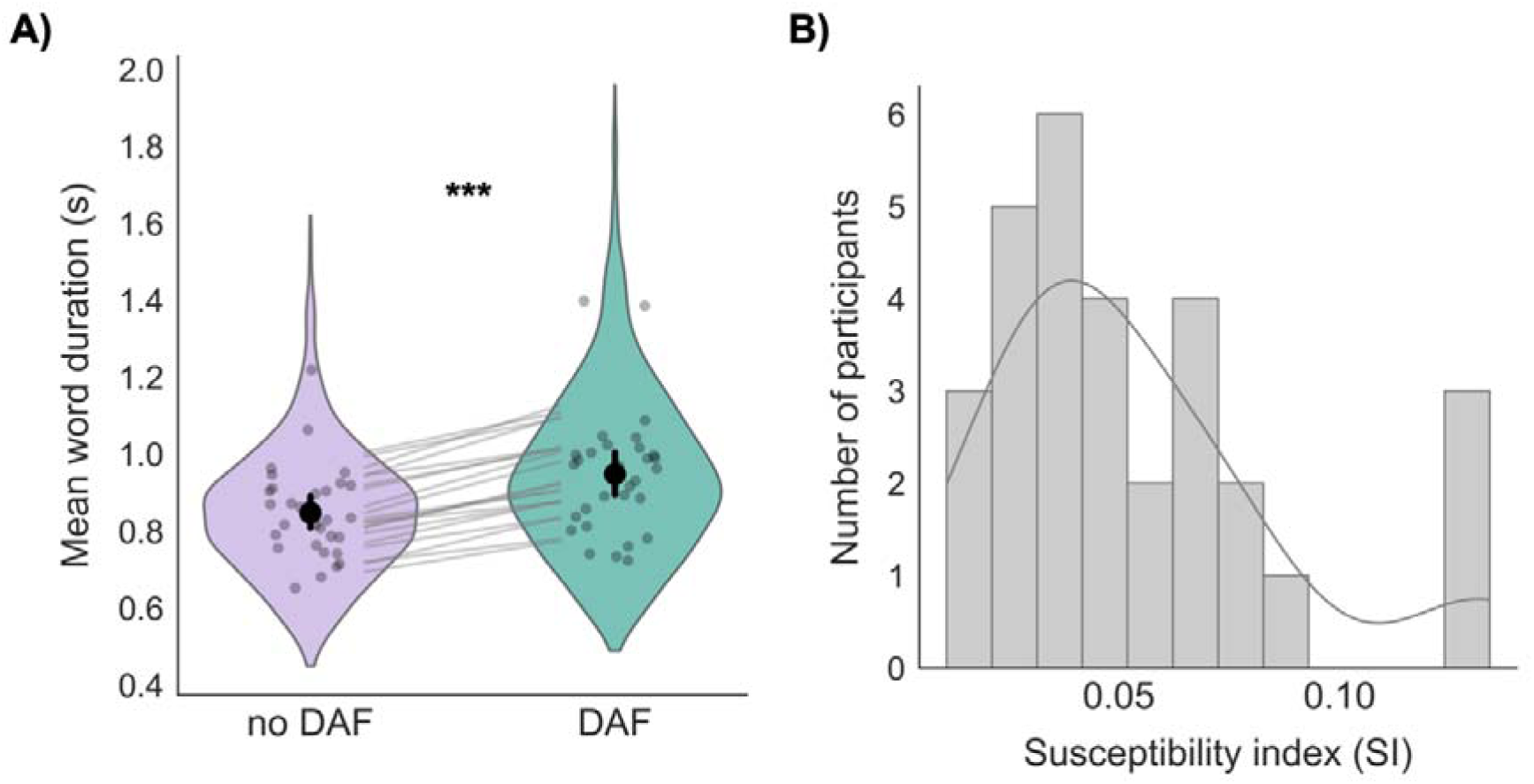
Delayed auditory feedback (DAF) slows speech. **(A)** Violin plots illustrate the distribution of word durations across all trials for the no-DAF (purple) and 200ms DAF (green) conditions. DAF significantly lengthened word duration (paired t-test: t = 7.859, p < 0.001). Gray pairing lines represent the mean change in duration for individual words, highlighting the consistent lengthening effect across the lexicon. Gray dots denote individual subjects (N=30). The central black markers and vertical error bars represent the group-level mean and 95% bootstrap confidence intervals, respectively. **(B)** The histogram and the overlaid density curve illustrate the distribution of the Susceptibility Index (SI) across all participants. Higher SI values indicate a greater degree of speech slowing under DAF relative to no-DAF condition.

Individual inspection of participants revealed substantial variability in susceptibility to DAF: some participants showed pronounced slowing, whereas others largely maintained their speech rate. To quantify this effect, a susceptibility index (SI) was computed for each participant by comparing word duration between the DAF and no-DAF conditions (see *Methods* for SI calculation). SI values varied widely across participants, ranging from 0.011 to 0.131 (**Fig. 1B, Fig. S1)**.

To assess whether DAF affected vocal pitch or intensity, we conducted a secondary acoustic analysis of the produced speech. Mean pitch and intensity values were extracted on a trial-by-trial basis using automated scripts in Praat (51). To account for inherent vocal differences between male and female speakers, values were standardized (z-scored) within gender prior to statistical analysis, thereby removing baseline gender-related differences while preserving within-subject variability. At the group level, DAF did not significantly modulate either acoustic measure. Standardized pitch values were comparable between conditions (paired *t*-test: *t* = −1.08, *p* = 0.29), with similar mean values observed for both males (no-DAF: 118.52 ± 15.32 Hz; DAF: 118.61 ± 16.05 Hz) and females (no-DAF: 207.82 ± 28.45 Hz; DAF: 206.54 ± 27.90 Hz). Likewise, vocal intensity did not differ significantly between conditions (*t* = −0.38, *p* = 0.70), again showing comparable values for males (no-DAF: 74.65 ± 9.45 dB; DAF: 75.02 ± 9.12 dB) and females (no-DAF: 76.12 ± 8.22 dB; DAF: 75.85 ± 7.95 dB).

### Group-Level Functional Activation for DAF

Speech production under delayed auditory feedback (DAF > no-DAF) elicited bilateral activation across speech-motor regions, with a more extensive recruitment observed in the right hemisphere (**Table S1, Fig. 2A–B**). To formally quantify this observation, we performed a hemispheric lateralization analysis at a statistical threshold of T > 2.75 (p < 0.01, two-tailed). At this level, all 31 subjects exhibited supra-threshold voxels and were included in the analysis, yielding a group mean Laterality Index (LI) of 0.198 (SD = 0.320, see **Fig. S2** for the distribution of LI values across participants). A one-sample t-test confirmed that this rightward shift was significantly greater than zero (t (30) = 3.448, p = 0.0017), representing a medium-to-large effect size (Cohen’s d = 0.619). The reliability of this lateralization was further validated by a bootstrapping procedure (10,000 resamples), which produced a 95% confidence interval [0.088, 0.306] that was entirely positive. This highly significant bootstrap p-value (p < 0.0001) suggests that right-hemisphere dominance is a robust and consistent feature of the neural response to DAF. Furthermore, sensitivity analyses conducted across a range of voxel-inclusion thresholds (T = 2.0 to 3.6, equivalent to p < 0.05 to 0.00) demonstrated that this rightward dominance remained stable regardless of statistical stringency. Across all tested levels, the bootstrap p-value remained highly significant (p < 0.0001), confirming that the right hemisphere is preferentially recruited for speech monitoring when auditory feedback is disrupted.

**Fig. 2.**
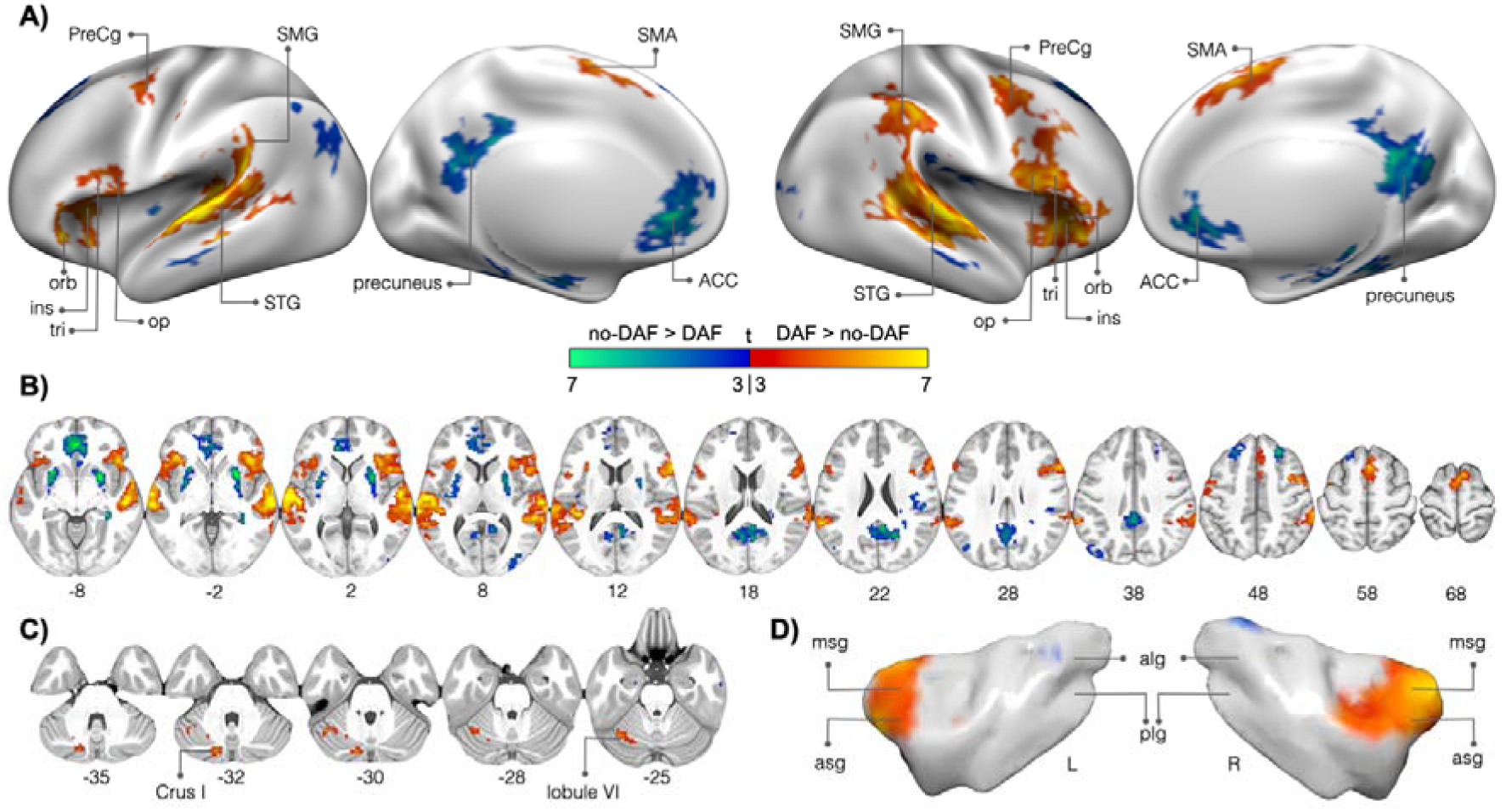
Whole-brain activations during speech production with delayed auditory feedback (DAF). **(A)** Cortical surface renderings (MNI152 inflated surface) showing group-level (N=31) activation for the DAF > no-DAF contrast (warm colors) and no-DAF > DAF contrast (cool colors) on left and right lateral and medial views. **(B)** Axial sections showing the same effects across the whole brain **(C)** Cerebellar sections **(D)** Left and right insular surfaces (Faillenot et al., 2017). Significant clusters identified in SPM12 (p < 0.001 voxel-level; p < 0.05 cluster-level FWE-corrected). All panels use the same color scale. SMA, supplementary motor area; PreCG, precentral gyrus; STG, superior temporal gyrus; SMG, supramarginal gyrus; IFG op/tri/orb, inferior frontal gyrus pars opercularis/triangularis/orbitalis; ACC, anterior cingulate cortex; ins, insula. asg (anterior short gyrus), msg (middle short gyrus), alg (anterior long gyrus), plg (posterior long gyrus).

Significantly greater activation for DAF was observed in the bilateral superior temporal gyri (STG) extending into the supramarginal gyrus (SMG), consistent with increased auditory and somatosensory processing demands. Enhanced responses were also found in speech motor regions, including the right inferior frontal gyrus (IFG), right supplementary motor area (SMA), and bilateral precentral gyri (PreCG), reflecting heightened demands on motor planning and articulatory control under perturbed feedback conditions.

Additional activation was detected in the left superior cerebellar regions, including Crus I and lobule VI (**Fig. 2C**), which have been previously shown to engage during both overt and covert speech production and are considered integral to the online control of articulatory movement sequences (52). Significant activation was also observed in the insula, most prominently in the anterior portion of the right insular gyrus (anterior and middle short gyri) (**Fig. 2D**). This portion of the insula has been consistently implicated in overt speech production (53), and damage to it is known to cause impairments in speech and orofacial motor control (54, 55).

The reverse contrast (no-DAF > DAF) revealed regions more active when speech unfolds with natural, unperturbed feedback. This analysis identified a frontal-parietal/medial-temporal network encompassing the left anterior cingulate cortex (ACC), right precuneus/posterior medial cortex, left angular gyrus, and bilateral hippocampal formation (**Table S1, Fig. 2A-B**). Increased activity was also observed in the bilateral putamen, along with smaller clusters (< 75 voxels) in the right Rolandic operculum, bilateral middle temporal cortices, left fusiform gyrus, right middle occipital gyrus, and right insula.

### Functional Associates of DAF Susceptibility

To investigate how brain activation patterns varied with individual susceptibility to DAF, we incorporated susceptibility indices (SI) as covariates in the group-level GLM analysis. This approach revealed a predominantly left-lateralized activation pattern (**Table S2, Fig. 3A-B**). Higher DAF susceptibility was associated with larger responses in the left inferior frontal gyrus (IFG), dorsal precentral gyrus (PreCG), supramarginal gyrus (SMG), cingulate cortex, and middle frontal gyrus (MFG), accompanied by strong engagement of the cerebellar vermis (lobules 4–5) (**Fig. 3C**). Significant clusters were also observed in the right hemisphere, including the supplementary motor area (SMA), IFG, and cerebellar lobule VI. In the left hemisphere, IFG activation extended into the anterior insula (middle short gyri) (**Fig. 3D**). For the reverse contrast (no-DAF > DAF), SI values were positively correlated with activity in the left angular gyrus.

**Fig. 3.**
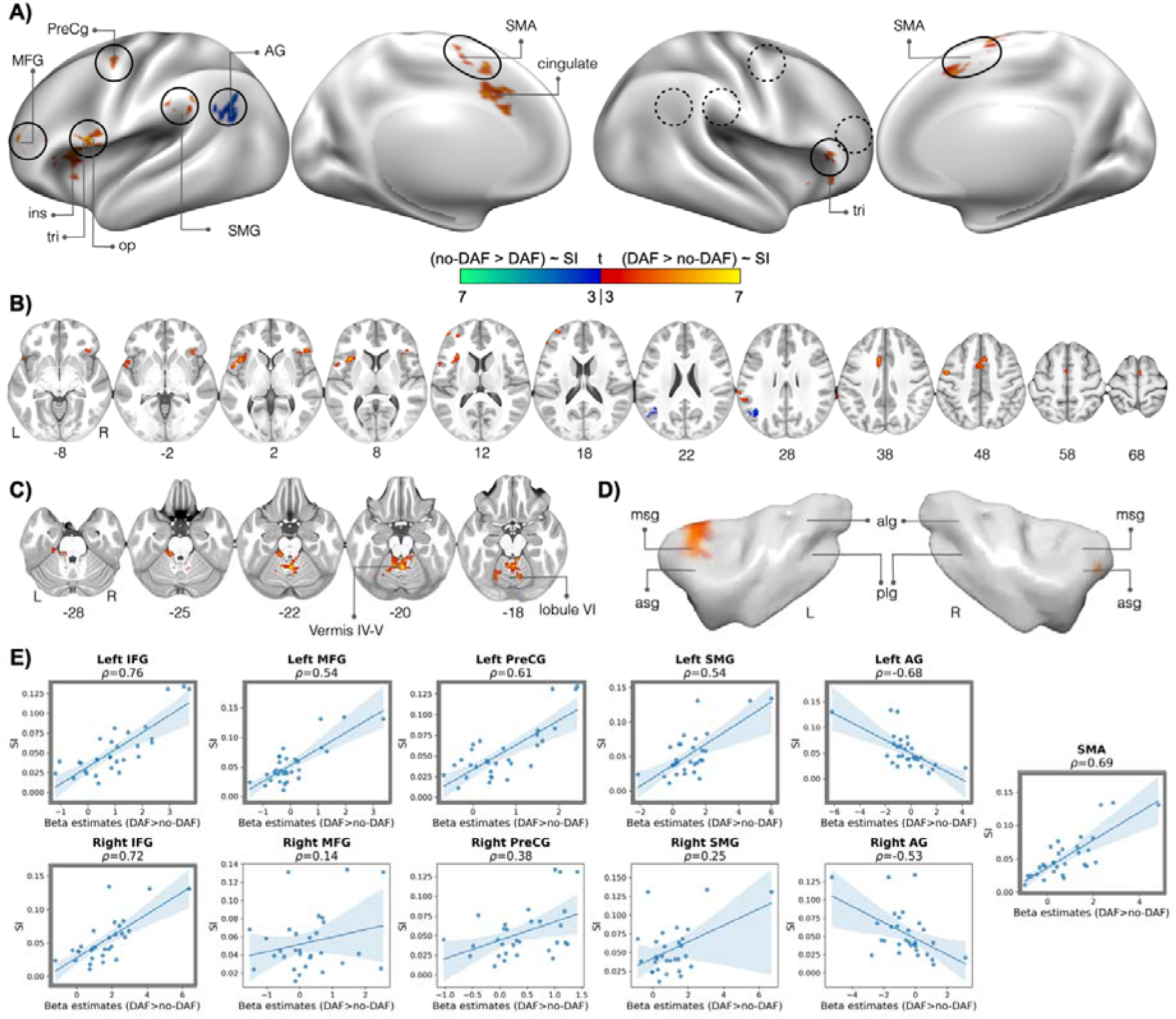
Whole-brain activations associated with individual susceptibility to delayed auditory feedback (DAF). **(A)** Cortical surface renderings (MNI152 inflated surface) showing regions where group-level activation (N=30) for the DAF > no-DAF contrast (warm colors) and no-DAF > DAF contrast (cool colors) significantly covaried with individual susceptibility indices (SI). Results are displayed on left and right lateral and medial views. Black outlines indicate the functional ROIs used for the analyses in Panel **E**; solid lines denote ROIs where significant correlations were observed in the whole-brain covariate analysis, and dashed lines represent their right-hemisphere homologues. **(B)** Axial sections showing the same effects across the whole brain **(C)** Cerebellar sections **(D)** Left and right insular surfaces (Faillenot et al., 2017). Significant clusters identified in SPM12 (p < 0.001 voxel-level; p < 0.05 cluster-level FWE-corrected). All panels use the same color scale. **(E)** Associations between SI and functional ROI activations; points are participants, lines show fitted trend with 95% CI, and Spearman rho values are shown on top of the plots. Scatter plots that are shown in a darker gray frame correspond to the functional ROIs indicated with solid black outlines in Panel **A**; A single ROI was defined for the left and right SMA. SMA, supplementary motor area; PreCG, precentral gyrus; STG, superior temporal gyrus; SMG, supramarginal gyrus; AG, angular gyrus; IFG op/tri/orb, inferior frontal gyrus pars opercularis/triangularis/orbitalis; MFG, middle frontal gyrus; ins, insula. asg (anterior short gyrus), msg (middle short gyrus), alg (anterior long gyrus), plg (posterior long gyrus).

To further illustrate the relationship between activation levels and SI values, we conducted a post-hoc functional ROI analysis based on the significant clusters identified in the group-level covariate analysis (see Methods). Specifically, ROIs were generated from clusters in the left IFG, MFG, PreCG, and SMG. In the right hemisphere, an ROI was generated from the activation cluster observed in the IFG. To investigate the laterality of these effects, we created homologous right-hemisphere ROIs for the MFG, PreCG, and SMG by mirroring the left-hemisphere clusters across the sagittal midline, as no significant clusters were observed in these regions at the whole-brain level. Additionally, the SMA cluster, which spanned both hemispheres, was treated as a single bilateral ROI, resulting in a total of 11 functional ROIs.

Mean beta estimates (DAF > no-DAF) were extracted from these functional ROIs to visualize the correlation of neural activation with participants’ SI values (**Fig. 3E**). In the left hemisphere, all ROIs exhibited a positive association with SI (Spearman’s rho > 0.5), except for the angular gyrus (AG), which showed a negative association (Spearman’s rho = −0.68). While these patterns are consistent with the whole-brain covariate analysis, no further inferential statistics were performed on these ROIs to avoid circularity. Notably, the homologous right-hemisphere AG ROI also displayed a negative association with SI values (Spearman’s rho = −0.53). Although this cluster did not emerge in the initial whole-brain analysis likely due to stringent cluster-extent thresholding, the increased sensitivity of the ROI-based approach revealed a robust relationship between right AG activity and individual susceptibility.

Overall, the activation patterns that scale with individual susceptibility exhibit a lateralization opposite to the average group-level effect observed for DAF > no-DAF, which showed stronger right-hemisphere activity (**Fig. 2**). These results indicate that participants who were more behaviorally affected by DAF show greater activation in left hemisphere speech motor regions, along with cerebellar areas.

To dissociate neural responses related to feedback monitoring from those driven by prolonged motor execution, we conducted an additional first-level parametric modulation (PMod) analysis in which trial-by-trial word duration was included as a regressor (see Supplementary Methods). This approach allowed us to account for variance associated with speaking duration and isolate neural responses specifically related to the sensory-motor mismatch induced by DAF. Importantly, the main clusters identified in the primary analysis (**Table S1; Fig. 2A-B**), including STG, IFG, SMG, and right preCG, remained significant after controlling for duration (**Fig. S3A; Table S3**). Conversely, activation in the left PreCG and the cerebellum was no longer significant after controlling for word duration, suggesting that activity in these specific regions is primarily attributable to the increased time required for speech production. Furthermore, when these duration-corrected contrasts were entered into a second-level analysis with participants’ SI as a covariate, significant correlations persisted in left-hemisphere speech regions (**Fig. S3B; Table S4**), indicating that the increased neural recruitment in susceptible individuals is driven by feedback mismatch, independent of the effects of prolonged speech duration.

### Structural Associates of DAF Susceptibility

Motivated by the perisylvian *f*MRI activation patterns in response to DAF (**Fig. 2A-B**) and the putative role of arcuate pathways in sensory-motor integration (41), we tested whether interindividual anatomical variation in the arcuate fasciculus (AF) segments relates to DAF susceptibility. We used subject-specific tractography to reconstruct the anterior (frontal-parietal), posterior (temporal–parietal), and long (temporal–frontal) arcuate segments in each hemisphere (**Fig. 4A-B**). Functionally, the cortical terminations of these segments correspond to motor-somatosensory (anterior), auditory-somatosensory (posterior), and auditory-motor (long) speech processing regions (56). All three segments were dissected in both hemispheres for every participant. For each tract, volume and hindrance-modulated orientational anisotropy (HMOA) (57) were extracted and correlated with individual

**Fig. 4.**
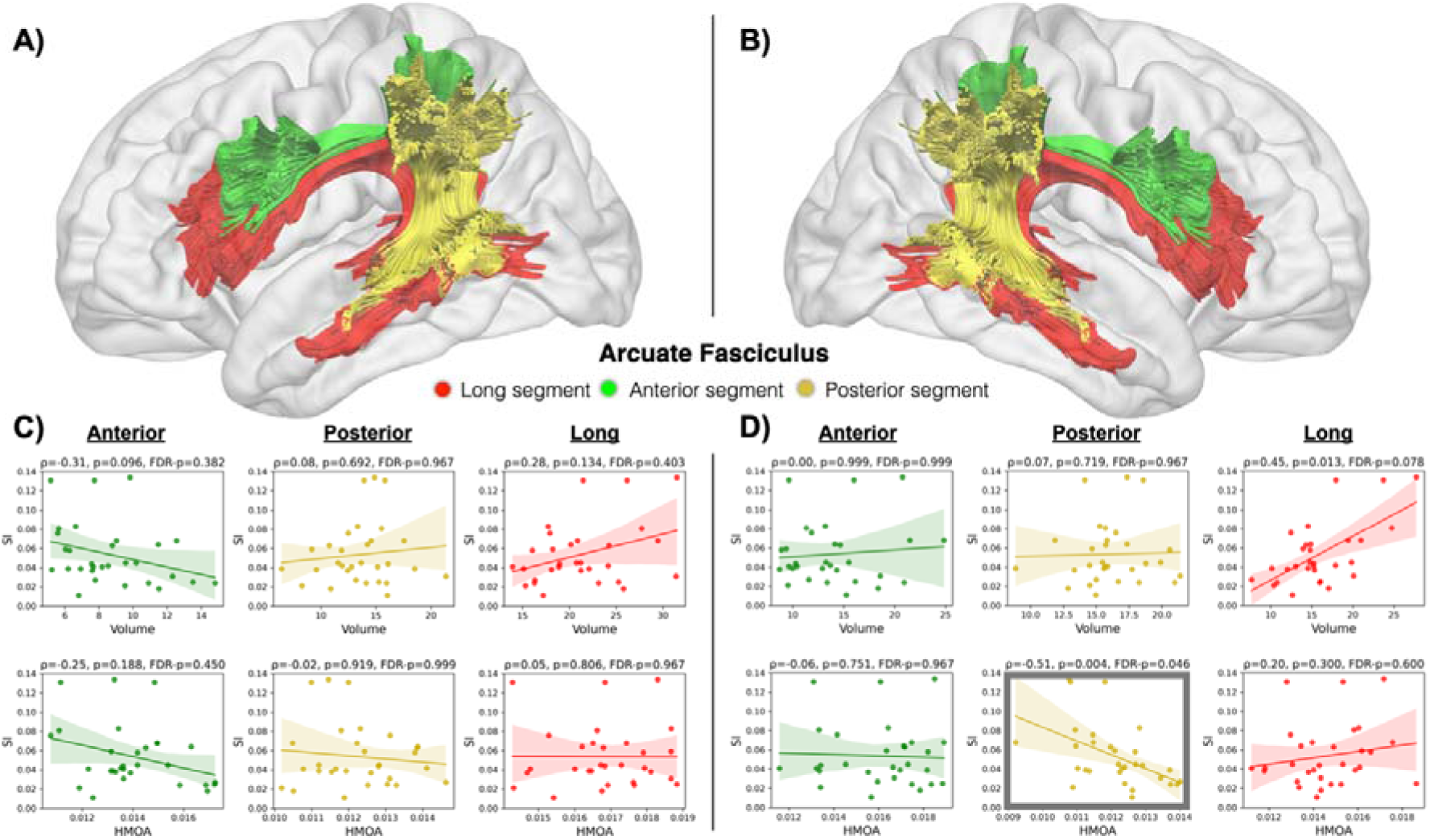
Correlations between individual susceptibility to DAF and the structural integrity of the three arcuate fasciculus segments. **(A-B)** Tractography-based dissection of the arcuate fasciculus (AF) into three segments: anterior (fronto-parietal), posterior (parietal-temporal), and long (fronto-temporal) **(C)** Left hemisphere: Associations (Spearman correlations) between SI and AF segment volume and HMOA; points are participants (N=30), lines show fitted trend with 95% CI **(D)** Right hemisphere: matching analyses for the homologous segments. Significant negative correlation was found for right posterior segment (scatter plot is shown in a darker gray frame). Spearman rho, p-value and FDR corrected p-values are shown on top of the plots. HMOA, hindrance-modulated orientational anisotropy; SI: Susceptibility index. susceptibility indices.

DAF susceptibility was negatively associated with HMOA of the right posterior segment (Spearman correlation: p = 0.004, FDR-p = 0.046), indicating that greater fiber density between posterior temporal and inferior parietal territories, connecting auditory and somatosensory speech region, was linked to a reduced behavioral impact of DAF. A positive association was observed between SI and volume of the right long segment (p = 0.013), connecting auditory and motor speech regions, however this association did not survive FDR correction (FDR-p = 0.078). No significant associations were observed for the right anterior segment or for any left hemisphere AF segments (**Fig. 4C-D**). These results point specifically to the right temporal-parietal arcuate as a structural support against DAF-induced dysfluency. For completeness, we also surveyed other canonical language-relevant white-matter pathways (e.g., FAT: frontal aslant tract and FIT: frontal-insular tracts), but none of these tract-behavior correlations survived multiple comparison correction (**Fig. S4 and S5**).

## DISCUSSION

Fluent speech depends on the alignment between internally predicted and externally perceived sensory feedback. When this alignment is perturbed, as under delayed auditory feedback (DAF), the brain must reconcile conflicting sensory inputs to maintain fluency. By linking behavioral susceptibility with both functional activation and white-matter connectivity, our results show that individual differences in this compensatory process reflect underlying neural architecture: DAF recruits a predominantly right-lateralized speech monitoring network, in which susceptibility is characterized by an increased recruitment of left-hemisphere monitoring homologues and a structural bias toward auditory-motor coupling. In contrast, resilience is linked to stronger structural connectivity supporting auditory-somatosensory integration.

Speech production models commonly describe sensorimotor interactions as a feedforward-feedback control system (38, 39). Within this framework, when a motor command is issued, an efference copy generates a neural prediction of the expected sensory consequences. This internal prediction is compared with incoming sensory input, enabling online correction of any discrepancies. Mismatch errors occur when the perceived sensory feedback diverges from the efference-based prediction. Such mismatches are detected by auditory comparator mechanisms in the posterior superior temporal gyrus (pSTG), as well as somatosensory comparators in the supramarginal gyrus (SMG). Corrective signals are transmitted to the inferior frontal gyrus (IFG), where articulatory plans are adjusted in real-time and relayed to the motor cortex, which subsequently sends articulatory commands to the vocal tract. This parallel system operating for auditory and somatosensory feedback (e.g., tactile sensations from the mouth and proprioception from the articulators) is thought to create a dual-control mechanism for speech production.

Our functional activation results converge on these described nodes of the feedforward-feedback loop. Consistent with a heightened demand on sensory processing and motor control in response to altered auditory feedback, DAF elicited activation in the STG, SMG, IFG and precentral gyrus (PreCG) (**Fig. 2**), predominantly in the right hemisphere.

Specifically, the dorsal PreCG activation aligns with our earlier electrocorticography findings in epilepsy patients, which showed selective engagement of this region during sentence production under DAF, supporting its role in maintaining speech fluency when dynamic auditory feedback processing is required to produce syllable sequences (29).

Beyond these core regions, the right hemisphere showed significant activation in the anterior portion of the insular gyrus, which is structurally connected with the IFG via frontal-insular tracts (58–60). Consistent with its proposed role as a relay between speech planning in IFG and motor execution in subcortical and cortical speech circuits (61), anterior insula activation may reflect the translation of phonetic intentions into coordinated vocal tract movements. Similarly, the SMG is thought to act as a key relay in the dorsal speech stream, supporting sensory-motor transformations by mapping phonetic representations onto articulatory movements (62, 63). These activations likely represent a functional network for integrating phonetic planning with sensorimotor control, enabling the rapid error monitoring and subsequent motor adjustments under DAF.

We also observed engagement of the right supplementary motor area (SMA) together with left cerebellar regions, which are structurally connected via crossed cortico-ponto-cerebellar and cerebello-thalamo-cortical pathways (64). Moreover, the SMA is reciprocally connected with the basal ganglia via thalamocortical loops and linked to the IFG via the frontal aslant tract (FAT) (65), forming a circuit that supports the initiation, sequencing, and timing of speech motor commands (66, 67). Consistent with these crossed pathways, co-activation of the right SMA with left cerebellar structures during DAF suggests increased demands on the temporal precision and coordination of articulatory movements. Given that DAF inherently slows speech, we carefully isolated the effect of increased speaking duration on these neural responses. Using a parametric modulation approach to account for trial-by-trial word duration, we found that key monitoring regions, including the STG, IFG, and SMG, remained robustly engaged (**Fig. S3A**), suggesting their activity specifically reflects the processing of feedback mismatch rather than extended motor execution. Interestingly, while these monitoring hubs remained significant, activation in the left PreCG and the cerebellum was largely accounted for by speaking duration, suggesting these specific nodes scale primarily with motoric workload.

Examination of the reverse contrast (no-DAF > DAF) highlighted a network of regions preferentially engaged during fluent speech under natural, unperturbed feedback conditions, revealing a distributed frontoparietal and medial-temporal network encompassing the left anterior cingulate cortex (ACC), right precuneus/posterior medial cortex, left angular gyrus, and bilateral hippocampal formation. This activation pattern overlaps substantially with the canonical default mode network (68), suggesting that when speech proceeds with intact feedback, the reduced demand for external monitoring may allow for a shift toward internally oriented processing. In this context, DMN engagement likely reflects the efficient, automated execution of speech supported by stable internal models. Additionally, increased activity was observed in the bilateral putamen. Given that the putamen is associated with motor sequence learning (69) and strongly responds to successful movement outcomes (70, 71), its increased activation for the no-DAF condition likely reflects the reinforcement of successfully produced articulatory sequences when speech outcomes align with internal predictions.

In contrast to the right-hemisphere bias observed at the group level in response to DAF, incorporating individual DAF susceptibility indices revealed a distinct shift in activation patterns (**Fig. 3**). Participants who were more susceptible to DAF showed stronger activation in several left-hemisphere speech motor homologues, including the IFG, SMG, SMA and PreCG. Critically, this correlation between susceptibility and left-hemisphere recruitment remained significant even after controlling for speaking duration (**Fig. S3B**), indicating that increased recruitment in more susceptible participants is not simply a byproduct of speaking longer. From a neural efficiency perspective (72, 73), the increased activation in susceptible individuals likely reflects greater resource recruitment to maintain motor control under conditions of high interference. In addition, greater DAF susceptibility was associated with heightened activity also in the cingulate cortex and MFG, the latter corresponding to the dorsolateral prefrontal cortex (DLPFC). These regions are commonly implicated in performance monitoring, cognitive control, and the allocation of executive resources during challenging tasks (74–76) Their engagement in more susceptible individuals may therefore reflect increased involvement of domain-general control mechanisms when speech production is disrupted by altered auditory feedback. These results differ from those reported by Agnew et al. (2018) (28), who found that resilience to DAF, measured by perceptual ratings of speech naturalness, was associated with increased right-hemisphere engagement in the putamen, ventral sensorimotor cortex, insula, and parietal operculum.

Although the two findings may appear to diverge, they likely reflect different aspects of the speech control system. Whereas the former study emphasizes neural processes supporting the maintenance of perceptual speech quality during sentence production, our study captures the degree of temporal disruption and compensatory effort required to sustain fluent speech under altered feedback.

The only region we observed to show increased activation in more resilient individuals was the bilateral angular gyrus (AG). The AG is often described as a cross-modal hub that integrates information from multiple sensory modalities (77, 78). It has also been implicated in detecting intersensory mismatches during action-feedback monitoring. In a paradigm conceptually similar to our DAF task, but introducing delays between somatosensory and visual signals, van Kemenade et al. (79) manipulated temporal delays between hand movements and their visual feedback during both active and passive movements. Using fMRI, they showed that activity in the AG correlated with participants’ ability to detect these intersensory mismatches, suggesting that this region monitors discrepancies between sensory signals. Our results extend these findings from hand movements to speech production and suggest that the AG, together with the SMG forming the inferior parietal lobule (IPL), constitutes a critical node for monitoring conflicts both across different sensory modalities and between external somatosensory feedback and internal motor predictions.

Given that speech monitoring relies on rapid information exchange across auditory, somatosensory and motor speech regions, we next asked whether interindividual differences in the white matter linking them could further explain variability in DAF susceptibility. We reconstructed the arcuate fasciculus in each hemisphere and examined its three components (**Fig. 4**): the long temporo-frontal segment, the posterior parieto-temporal segment, and the anterior fronto-parietal segment (41), whose cortical terminations overlapped with our functional cluster peaks in IFG, SMG and pSTG (56). Across participants, susceptibility to DAF increased with the volume of the right long segment (uncorrected p = 0.013, FDR-p = 0.078). Although this trend did not survive multiple comparison correction, it is anatomically coherent (**Fig. 5**): a more substantial right temporo-frontal conduit would be expected to up-weight auditory input to frontal speech motor regions; when feedback is delayed, that heavier weighting manifests as greater behavioral slowing. This implies that individuals who are more affected by DAF may rely more heavily on auditory input, making them more vulnerable when auditory feedback is disrupted.

**Fig. 5.**
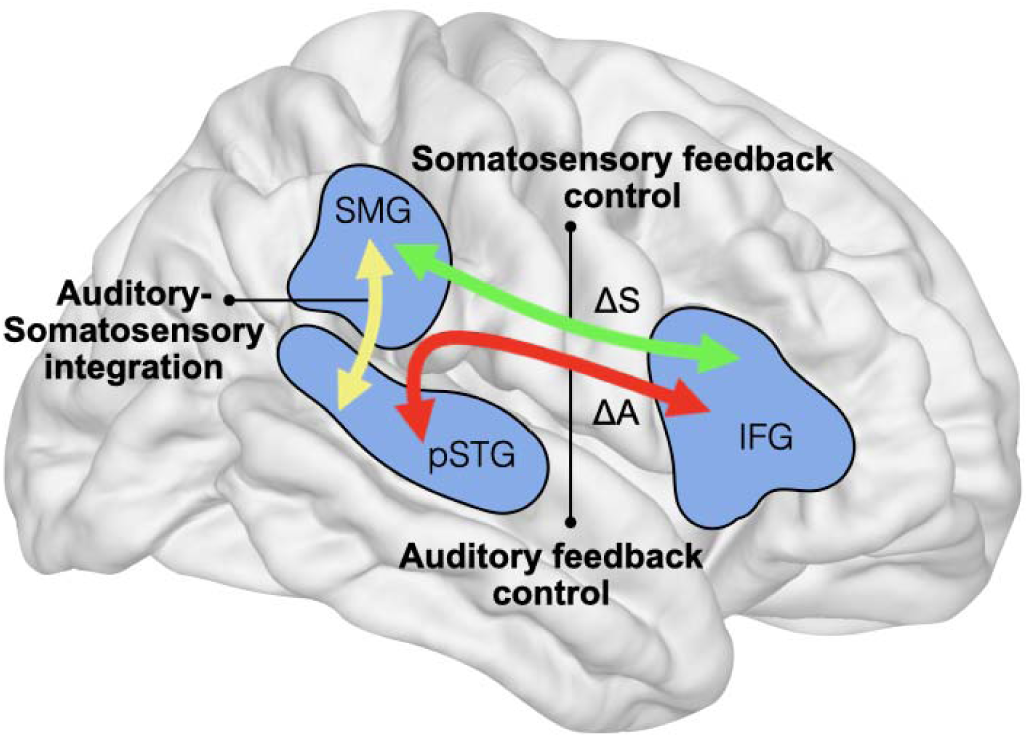
Schematic synthesis of functional and structural correlates of DAF processing. The posterior superior temporal gyrus (pSTG) and supramarginal gyrus (SMG) serve as key nodes for encoding auditory and somatosensory corrective signals (ΔA and ΔS), respectively. These signals are then relayed to speech motor planning regions like the inferior frontal gyrus (IFG) to adjust the ongoing motor commands. The crosstalk between the pSTG and SMG enables an efficient weighing and integration of the two sensory feedback streams for motor control. The functional connections that enable this monitoring process are supported by the structural pathways of the arcuate fasciculus (AF) depicted by the double-sided arrows highlighting anterior (green), posterior (yellow) and long (red) segments of the AF.

Conversely, a significant negative correlation emerged between DAF susceptibility and fiber density (HMOA) of the right posterior AF segment (uncorrected p = 0.004; FDR-p = 0.046). This finding suggests that stronger connectivity between pSTG and SMG supports cross-sensory interactions and more effective weighting and integration of multiple feedback sources, enabling individuals to better maintain fluency when auditory and somatosensory channels provide conflicting information. DAF fundamentally disrupts sensorimotor processing by introducing a temporal misalignment between the motor commands and their auditory consequences, thereby desynchronizing auditory and somatosensory feedback streams. When the brain attempts to correct for a perturbation in auditory feedback, it may require deviating from its somatosensory targets, and vice versa. This inherent trade-off suggests that effective crosstalk and dynamic integration between auditory and somatosensory feedback systems are critical for maintaining speech fluency under feedback perturbation. These observations motivate a tract-specific account of feedback control, one in which the right long AF segment can amplify auditory-to-motor drive, whereas the right posterior AF segment provides a stabilizing substrate by supporting auditory-somatosensory integration.

Taken together, these functional and structural findings suggest that while DAF processing is predominantly right-lateralized at the group level, individual resilience is reflected in distinct neural profiles. Susceptible individuals more heavily recruit frontal motor-control regions (IFG, SMA) and left-hemisphere speech motor regions (SMG and preCG) to manage the increased demands of altered feedback, a pattern reflecting compensatory neural effort. In contrast, resilient individuals exhibit enhanced engagement of the bilateral AG alongside stronger structural connectivity in the right posterior arcuate fasciculus. This suggests that resilient individuals leverage a posterior “auditory-somatosensory integrator” circuit to efficiently resolve sensory conflicts, thereby reducing the need for the increased recruitment of speech motor control regions observed in more susceptible individuals. Consequently, resilience appears to be a function of a more focal and efficient right-lateralized network. Critically, our results build upon current speech models by demonstrating that the right hemisphere’s role extends beyond the processing of vocal pitch or formant manipulations to the maintenance of speech coordination and fluency.

These findings also raise new questions about how such feedback control mechanisms are engaged under different task contexts and levels of feedback predictability. While our randomized paradigm characterizes neural mechanisms of online speech monitoring, alternative designs could address complementary aspects of feedback control. For example, cued paradigms in which perturbations are anticipated may allow investigation of pre-stimulus sensory reweighting, whereby top-down mechanisms regulate the gain of auditory feedback relative to somatosensory inputs or internal predictions (80). In addition, blocked designs with consistent perturbations could be used to examine whether the susceptibility profiles identified here predicts the rate of sensorimotor adaptation, reflecting the brain’s ability to use persistent error signals to update feedforward motor commands across trials (81). Together, such approaches may help clarify whether resilience to altered auditory feedback reflects a relatively stable neural trait or a more dynamic, context-dependent state characterized by flexible regulation of feedback gain and the faster updating of motor plans.

In summary, by leveraging a fluency-perturbing manipulation (DAF), an individual behavioral susceptibility index, whole-brain *f*MRI, and tract-specific diffusion metrics, we provide converging functional and structural evidence that resilience or vulnerability to feedback disruption reflects differential recruitment of speech motor control networks and variability in the strength of auditory-somatosensory connections. These findings move the field beyond group averages to a mechanistic, person-specific account of speech monitoring, with translational implications for understanding fluency breakdowns in disorders such as stuttering and relevant speech technologies.

## LIMITATIONS

Despite these compelling findings, several features of our study design qualify our inferences and point to directions for future research. First, our focus on auditory perturbation precludes direct assessment of how auditory and somatosensory feedback are jointly weighted. Incorporating a somatosensory perturbation paradigm would enable a comprehensive investigation of this interaction (82, 83), while also allowing us to assess participants’ somatosensory susceptibilities.

Second, our behavioral index was based primarily on word duration. While this provided a continuous measure of susceptibility, richer fluency measures (e.g., pause structure, speech rate dynamics, error profiles, and articulatory kinematics) would allow for a more complete behavioral characterization of DAF susceptibility. Prior work in connected speech (e.g., sentences or paragraphs) has shown that, in addition to speech slowing, DAF can elicit dysfluencies such as syllable repetitions, misarticulations, or segmental errors (15, 84–86). Speech slowing reflects the real-time integration of DAF into the motor plan, whereby a detected mismatch triggers a compensatory lengthening of speech. In connected speech, however, over reliance on auditory feedback can lead to cascading error signals that cause the system to effectively “reset” the motor plan, manifesting as overt dysfluencies, as suggested by a computational model of stuttering (87). In single-word production tasks with DAF, dysfluencies are relatively rare compared to connected speech, as this design reduces opportunities for cascading errors while still engaging feedback-based monitoring and control processes. Although our use of a word production task, rather than longer connected speech segments, precluded the analysis of overt dysfluencies, it was a necessary design choice to mitigate motion and susceptibility artifacts in the scanner. This approach ensured high sensitivity in critical regions of interest, including the inferior frontal, anterior temporal, and cerebellar cortices, which are particularly susceptible to signal dropout during overt speech.

Finally, we acknowledge the inherent limitations of our imaging modalities. Our *f*MRI results are correlational and do not establish the directionality of information flow within the speech motor network. Model-based approaches (e.g., Dynamic Causal Modeling), brain perturbation (e.g., TMS), or intracranial recordings, would provide stronger causal leverage. Additionally, while diffusion tractography is affected by crossing fibers and partial-volume effects, we used a high diffusion weight and advanced spherical deconvolution to overcome these limitations to the best of our ability. Our a priori focus on the arcuate fasciculus increases the interpretability of our findings but risks overlooking other relevant pathways. Although exploratory analyses of other tracts (FAT and FIT 1-5, **Fig. S4 and S5**) did not survive multiple comparison correction, they suggest clear avenues for future hypothesis-generating work. These constraints outline a clear roadmap for the field: larger datasets with richer behavioral assays, the incorporation of somatosensory perturbations, causal and computational assays of network dynamics, and multimodal microstructure to further refine the structure-function account of feedback control.

## METHODS

### Participants

31 native Dutch speakers (18 females, age: 24 ± 6 years) were recruited through the Radboud SONA research participation system. All participants presenting without a history of hearing or language impairments were included in the study after providing written informed consent prior to the start of the experiments. The research was approved by the Ethics Committee of the Faculty of Social Sciences at Radboud University (ECSW-2020-046) and conducted in accordance with the Declaration of Helsinki.

### Experimental Procedure

The experiment consisted of three conditions: passive listening, speaking with immediate auditory feedback (no-DAF), and speaking with delayed auditory feedback (DAF). Passive listening condition was included as an auditory reference, allowing comparison between externally generated and self-generated speech during production (**Fig. S6**). During passive listening trials, participants were presented with the audio recordings of 20 different words spoken by a female native Dutch speaker. The words consisted of three syllables and had a mean duration of 0.72 seconds. Participants were instructed to look at the crosshair in the middle of the screen and passively listen to prerecorded words. For the two speaking conditions, the same list of words was presented on the screen as text, one word per trial. The words appeared as red text and participants were instructed to read them out loud as soon as the text turned green. During these speaking trials, auditory feedback was presented either simultaneously (no-DAF) or with a 200 ms delay (DAF) through insert earphones.

Each of the 20 words was repeated 2 times for each condition, which resulted in 40 trials per condition and a total of 120 trials for all three conditions. Trials for the three conditions were presented randomly in a jittered event-related *f*MRI design. To optimize the estimation of the Hemodynamic Response Function (HRF) and maximize statistical efficiency, null trials (fixation periods) were pseudo-randomly interspersed between experimental trials, ensuring a minimum inter-stimulus interval (ISI) of 3 seconds. During these null trials, participants passively fixated on a central crosshair. The randomization, including the optimal distribution of conditions and null intervals, was determined using *OptSeq* (88).

### Neuroimaging Data Acquisition

Structural (T1w, diffusion-weighted) and functional (*f*MRI) data were acquired on a 3T Siemens MAGNETOM Prisma Fit scanner using a 32-channel head coil at the Donders

Institute, Nijmegen. Structural data were acquired using a T1-weighted 3D magnetization-prepared rapid gradient-echo (MPRAGE) sequence (TR = 2300 ms, TE = 3.03 ms, TI = 1100 ms, flip angle = 8°, 192 sagittal slices, voxel size = 1 × 1 × 1 mm³, FOV = 256 × 256 mm², GRAPPA acceleration factor = 2). The total acquisition time was 5 minutes and 21 seconds.

Diffusion-weighted imaging was acquired as a single-shot spin-echo echo-planar imaging (EPI) acquisition with GRAPPA acceleration (factor = 2) and multi-band acceleration (factor = 3). Acquisition parameters were TR = 2940 ms, TE = 74.8 ms, flip angle = 90°, FOV = 216 × 216 mm², matrix = 120 × 120, isotropic voxel size of 1.8 mm³, 81 axial slices with interleaved acquisition, and anterior-to-posterior phase encoding. Diffusion gradients were applied along 86 non-collinear directions at a b-value of 1250 s/mm^2^ and 85 non-collinear directions at a b-value of 2500 s/mm^2^. Twelve interleaved volumes were collected without diffusion weighting (b0 volumes). Total scan time was approximately 10 minutes. Additionally, seven b0 volumes were acquired with the reverse phase encoding direction (posterior-to-anterior) to facilitate susceptibility distortion correction.

Functional data were acquired using a T2* weighted gradient-echo echo-planar imaging (EPI) sequence with a sparse sampling design (TR = 3000 ms, TA ≈ 1000 ms, TE = 33 ms, flip angle = 80°). Each volume consisted of 52 interleaved transverse slices (voxel size = 2.4 × 2.4 × 2.4 mm³, FOV = 210 × 210 mm²) acquired with a multiband acceleration factor of 4. This sparse-sampling approach was utilized to prevent the acoustic contamination of auditory stimuli by scanner noise, providing a 2-second silent window for speech production and perception between successive 1-second acquisition periods.

A total of 400 volumes were collected per run, consisting of 120 task-related volumes (40 per condition) and 280 null-trial volumes. The timing of each trial was strictly synchronized with the scanner’s trigger. At the offset of the scanner noise for each task volume, a 2 s silent window commenced. For ‘Listen’ trials, the auditory stimulus was initiated at this onset. For speaking trials (DAF and no-DAF), a dual-cue strategy was employed: the visual text was initially presented in red and, after 1 s, changed to green simultaneously with the cessation of scanner noise, signaling the participant to begin speaking. Participants were instructed to complete their vocal response within the silent gap. Given that the task required the production of single words, the 2 s window provided ample time to complete the utterance. This design ensured that the blood-oxygen-level-dependent (BOLD) signal was sampled near its peak, while avoiding acoustic contamination of the recordings. All functional data were collected in a single run lasting approximately 20 minutes.

Stimulus presentation was executed using Matlab Psychtoolbox-3 running on an HP Intel Core i7 laptop with a Windows 10 operating system. As participants read aloud the words, their voices were recorded using an MRI compatible microphone (Optoacoustics, FOMRI-III) and played back using a Focusrite Scarlett 2i2 audio interface and S14 MRI compatible insert earphone. For the DAF condition, the recorded audio was delayed using Matlab’s Audio Toolbox functions.

### DWI Analysis and Tractography

The preprocessing of the DWI data followed our in-house pipeline (89, 90). Briefly, DWI data were corrected for thermal noise (91) and Gibbs ringing artefacts (92) using tools implemented in *TORTOISE* (Irfanoglu et al., 2017). The pair of b0 images acquired with reverse phase encoding was used to estimate a geometric distortion field in *topup* (94). The tool *eddy* was then used to correct the full DWI series for head motion, eddy current distortions, slice-to-volume motion, signal dropout, and susceptibility distortions based on the *topup* field, including the effect of motion on these distortions (95–98). DWI data from the b = 2500 s/mm^2^ shell were used for tractography. The data were modelled in StarTrack (https://nbl-research.github.io) using the damped Richardson-Lucy spherical deconvolution algorithm (57, 99) and the following parameters: fiber response function _α_ = 1.5; number of iterations = 200; damping parameter _η_ = 0.0015; adaptive regularization parameter _ν_ = 16. Tractography was then performed using a multi-fiber Euler-like algorithm and the following parameters: minimum amplitude = 0.0025; angle threshold = 40°; step size = 0.9 mm; fiber length range = 20–300 mm. The anisotropic power (AP) maps (100) generated by StarTrack were used to normalize each participant’s DWI data and tractography to the common space of the ICBM 2009a nonlinear symmetric MNI template (101) using diffeomorphic registration in ANTs (Avants et al., 2008, 2014). MegaTrack (102) was used to perform semi-automatic virtual dissections using the region-of-interest placement described in Forkel et al. (103). The use of the symmetric template allowed us to use the symmetric flipping option in MegaTrack, which renders the operator blind to the hemisphere they are dissecting, hence removing any user bias. The interactive virtual dissections were performed in TrackVis (http://trackvis.org). Once the dissections were complete, tract volume and the hindrance modulated orientational anisotropy (HMOA) (57) were extracted in native space for each participant and each tract.

Tract volume (in milliliters, mL) was calculated by multiplying the volume of a single voxel (5.832×10-3 mL) by the total number of voxels visited by the tract. HMOA was extracted directly from the tractography data at each streamline point and was averaged over the entire tract. HMOA corresponds to the amplitude of the fiber orientation distribution (unitless) at each point in the tractogram and reflects the apparent fiber density at that point, among other measures of microstructural organization. For the main analysis, this resulted in three tracts (the three AF segments) in each hemisphere, with two metrics each (volume for macrostructure and HMOA for microstructure), resulting in a total of 12 variables that were tested for correlations with SI. The false discovery rate (FDR) method (104) was used to compensate for these multiple comparisons.

### *f*MRI Analysis

#### Preprocessing

*f*MRI preprocessing was performed using fMRIPrep 23.1.0 (105, 106). The T1-weighted images were spatially normalized to MNI standard space (MNI152NLin2009cAsym) after skull stripping and segmentation. The functional images were skull stripped, head motion parameters were estimated, followed by slice-time correction. The BOLD time-series were then co-registered to the T1w image and finally resampled to standard space (see *Supplementary Material* for a detailed description of the fMRIPrep preprocessing pipeline).

#### General Linear Model (GLM) analysis

First, we smoothed the preprocessed functional images in MNI space with an isotropic Gaussian kernel of 4 mm FWHM in SPM12 in Matlab R2023b. We then computed a general linear model in SPM12 with the following condition regressors: DAF, no-DAF, and Listen, including their temporal derivatives. In addition, we added head motion parameters estimated by fMRIPrep as confound regressors. The model was estimated after high-pass filtering and then we computed contrasts between the DAF and no*-*DAF conditions using weights [1 - 1]. The contrast images were entered into a group-level one-sample t-test, where we also included participant-specific susceptibility indices (see *Behavioural Analysis and Susceptibility Index (SI) Calculation* section) as a covariate. We then generated contrasts for the second-level model for the DAF and no-DAF difference and for the SI covariate. Statistical maps were thresholded at the voxel level (p < 0.001 uncorrected) and subjected to a cluster-level extent threshold (*p* < 0.05 family-wise error corrected at the cluster-level) to identify significant activation clusters.

#### Visualization of *f*MRI results

Thresholded group-level statistical maps were projected onto the cortical surface of the MNI152 template using in-house MATLAB scripts. The template surface was first reconstructed in FreeSurfer (https://surfer.nmr.mgh.harvard.edu), producing pial and white matter boundary surfaces for each hemisphere. For each surface vertex, voxel values were sampled using nearest-neighbor interpolation at nine equidistant points along the surface normal between the corresponding pial and white matter locations, and the maximum sampled value was assigned to that vertex. This approach balances sensitivity and specificity: sampling across cortical depth captures activation that may be offset from the surface due to partial-volume effects or minor registration errors, while taking the maximum rather than averaging preserves the peak statistical value at each cortical location without diluting it with surrounding non-activated voxels. The last point is especially important given the projection was applied to thresholded maps. The resulting surface overlays were displayed on the inflated surface representation in SurfIce (https://www.nitrc.org/projects/surfice). Voxel-based (slice) displays of activation maps were prepared using MRIcron (https://www.nitrc.org/projects/mricron).

#### Lateralization analysis

To quantify the hemispheric lateralization of the *f*MRI response for the DAF and no-DAF difference, a Laterality Index (LI) was calculated for each participant based on the distribution of significantly activated voxels. First, subject-specific T-statistical maps were thresholded at T > 2.75 (equivalent to p < 0.01, two-tailed). Using the image affine matrix, voxels were categorized into left and right hemispheres based on their MNI x-coordinates (x < 0 and x > 0, respectively). The LI was calculated for each participant using the standard formula: LI = (V_R_ - V_L_) / (V_R_ + V_L_), where V_R_ and V_L_ represent the count of supra-threshold voxels in the right and left hemispheres. Under this convention, a positive LI indicates right-hemisphere dominance, while a negative LI indicates left-hemisphere dominance. At the group level, the distribution of LI values was assessed using a one-sample t-test against a null hypothesis of zero (no lateralization), and effect sizes were calculated using Cohen’s d. To ensure the robustness of the group-level findings, a bootstrapping procedure was implemented (10,000 iterations with replacement). This was used to determine the 95% bias-corrected confidence intervals (CI) and a two-sided bootstrap p-value for the mean LI.

#### Functional Region of Interest (ROI) analysis

To visualize the relationship between SI and neural activation, we extracted mean parameter estimates (DAF > no-DAF) from clusters identified in the group-level SI covariate analysis. Significant clusters were defined based on the whole-brain group contrast thresholded at a voxel-wise level of p < 0.001, with cluster-extent FWE correction at p < 0.05. Mean activation values were extracted from these clusters using the ROI extraction tools implemented in MarsBaR in SPM12 (107). When a significant cluster was identified in one hemisphere but not the contralateral hemisphere, a homologous ROI was created by mirroring the cluster across the mid-sagittal plane. Specifically, the x-coordinates of all voxels within the original cluster were sign-flipped in MNI space to generate a contralateral homologue of identical size and shape. This procedure allowed for symmetric hemisphere-wise comparisons while preserving the functional definition of the ROI.

### Behavioural Analysis and Susceptibility Index (SI) Calculation

In-scanner voice recordings of the participants were analyzed using Praat (51). Due to a recording failure, one participant was excluded from all behavioral analyses, resulting in a final behavioral sample of N=30. For each trial, word onset and offset were manually marked on the audio waveform and spectrogram to determine word duration. These boundaries were then saved as time-aligned TextGrids for subsequent acoustic analysis. Acoustic analysis of the produced speech was conducted using custom *Praat* scripts to extract pitch (F0: fundamental frequency) and intensity on a trial-by-trial basis. For F0 estimation, an autocorrelation-based method was employed with gender-specific search ranges (Females: 100-500 Hz; Males: 75-300 Hz) and a 1-ms time step. To ensure data quality, octave-jump correction was applied to the resulting contours. Vocal intensity was calculated by generating intensity objects with a 100 Hz floor. Both measures were constrained to the onset and offset of each produced word using the time-aligned TextGrids. Mean F0 (in Hz) and mean intensity (in dB) were then calculated for each interval and exported for statistical analysis.

Paired t-tests were performed to test the effect of auditory feedback delay on word duration, vocal pitch, and intensity by comparing the DAF and no-DAF conditions across participants.

To quantify individual susceptibility to DAF, a susceptibility index (SI) was computed for each participant comparing word durations for the different conditions using the formula:

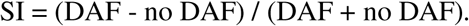

This index captures the degree to which speech slowing under DAF affects each individual. The SI ranged from −1 to 1, with higher values indicating greater susceptibility to DAF.

## Acknowledgements

The authors thank José Marques for programming the MRI acquisition protocol and Marteen van den Heuvel for programming the stimulus presentation. We also thank the members of the Neurobiology of Language department at the Max Planck Institute for Psycholinguistics and the Clinical Neuroanatomy of Language research group at the Donders Institute for their valuable feedback throughout this project.

## Data Availability

Preprocessed fMRI data will be made available via the Radboud Data Repository (https://data.ru.nl/). Group-level statistical T-maps and correlational spreadsheets linking individual tract metrics to susceptibility indices will be shared on GitHub (https://github.com/MugeOzker).

## Funding

This work was supported by the Dutch Research Council NWO Language in Interaction Grant (024.001.006), Donders Mohrmann Fellowship on ‘Neurovariability’ No. 2401515 (SJF), the Dutch Research Council NWO Aspasia Grant ‘Human individuality: phenotypes, cognition, and brain disorders’ (SJF) and the NWO SSH Open Competition grant (SJF).

## Notes

### Competing Interest Statement

The authors have declared no competing interest.

### Summary of Updates

We have updated the manuscript with new lateralization, functional ROI, and voice acoustic analyses. Additionally, we implemented a parametric modulation analysis to account for speaking duration. The figures, introduction, and discussion have been extended to reflect these additional findings.

