## Supplementary Information for "Individual Differences in Speech Monitoring: Functional and Structural Correlates of Delayed Auditory Feedback"

**SUPPLEMENTARY MATERIAL**

**Supplementary Table 1:** Significant fMRI clusters (voxel-level p < 0.001; cluster-level p < 0.05, FWE-corrected) showing greater activation for delayed auditory feedback (DAF > no-DAF) and for immediate feedback (no-DAF > DAF) during speech production.

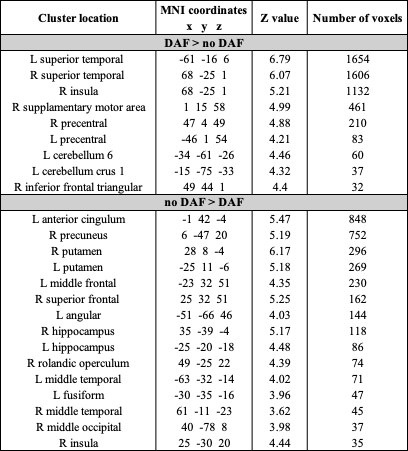

**Supplementary Table 2:** Significant fMRI clusters (voxel-level p < 0.001; cluster-level p < 0.05, FWE-corrected) showing correlations between neural activation (DAF > no-DAF and no-DAF > DAF) and individual susceptibility to DAF (SI).

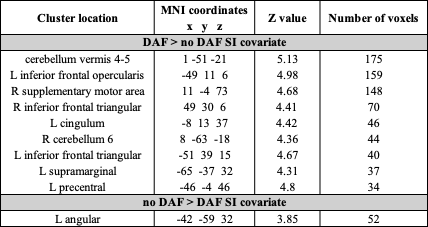

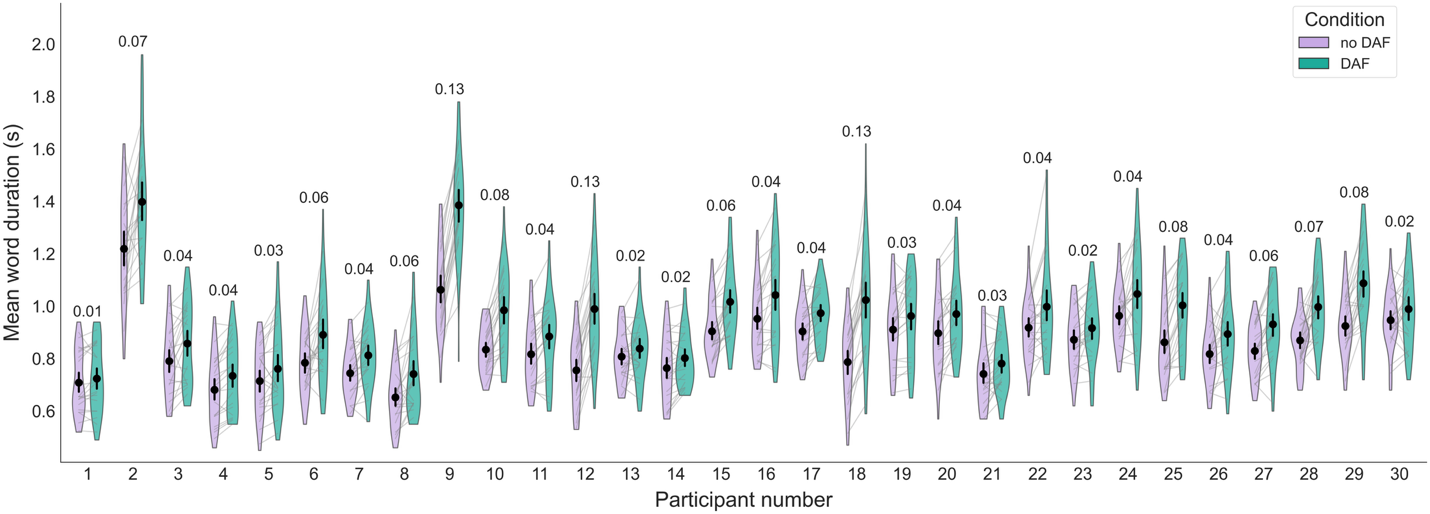

***Supplementary Figure 1:*** *Violin plots illustrate the distributions of word durations for each participant under no-DAF (purple) and DAF (green) conditions. Gray pairing lines represent the mean change in duration for individual words. Black markers and vertical error bars denote the participant-specific mean and 95% bootstrap confidence intervals, respectively. The numerical value above each participant’s plots indicates their Susceptibility Index (SI), representing the standardized magnitude of speech slowing under DAF. While across the sample (N=30), a consistent increase in word duration is observed during DAF trials, the degree of interference varies significantly between individuals as reflected by the SI values.*

### ***Supplementary Figure 2:*** ***Distribution of hemispheric functional lateralization for DAF processing****. Histogram of the Laterality Index (LI) across participants (N = 31), calculated from the number of voxels showing significant activation (T > 2.75) for the DAF > no-DAF contrast. Positive LI values indicate right-hemisphere dominance, while negative values indicate left-hemisphere dominance. The dashed black line (LI = 0) represents symmetrical bilateral activation, and the solid red line indicates the group mean. The distribution reveals a right-lateralized trend in the recruitment of the monitoring network, with a Kernel Density Estimate (KDE) overlay showing the underlying probability density of the participant sample.*

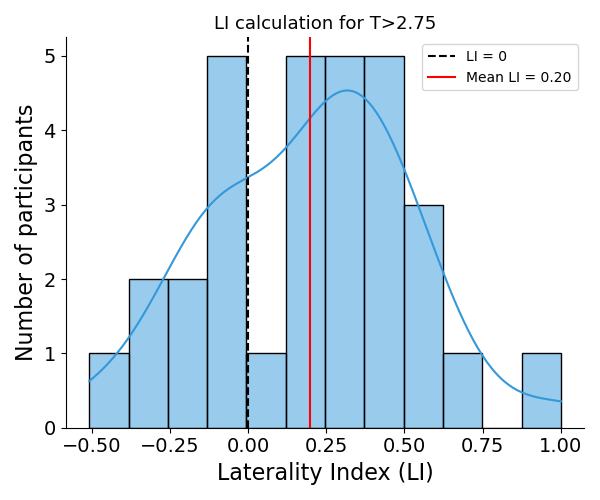

**Dissociating Feedback Monitoring from Motor Execution**

**Parametric modulation analysis:** To disentangle the neural response to feedback mismatch from the duration of motor execution, a supplemental control analysis was conducted at the first level. For each participant, a GLM was constructed in SPM12 that included a single "Speech" regressor containing the onsets of all speaking trials. Two Parametric Modulators (PMods) were entered in a hierarchical sequence: (1) Word Duration, representing the trial-by-trial length of the speech sample, and (2) Delay Condition, a binary vector representing the feedback type (DAF vs. no-DAF). The default Gram-Schmidt orthogonalization in SPM12 (orth = 1) was employed, which assigns all shared variance between the two modulators to the first entered variable (Word Duration). Consequently, the second PMod (Delay) captured only the unique, residual variance associated with the feedback mismatch that could not be explained by the length of the trial. The contrast images (DAF > no-DAF) generated from the first-level hierarchical PMod model were entered into random-effects group-level regression analyses.

Even after this conservative analysis, key regions in the STG, IFG, SMG, and right PreCG remained significantly active (p < 0.001 voxel-level; p < 0.05 cluster-level FWE-corrected; **Table S3, Figure S3A**). The persistence of these clusters indicates that these regions are specifically engaged in processing the sensory-motor mismatch inherent to DAF, independent of speech duration. In contrast, activation in the left PreCG and cerebellum was markedly reduced or absent compared to the activation identified prior to the PMod analysis (**Table S1, Figure 2A-B**). This dissociation suggests that whereas the monitoring network (STG/SMG/IFG) tracks feedback error, left primary motor cortex and cerebellar regions largely scale with motoric workload and trial duration.

To further examine whether these duration-corrected monitoring activity predict individual behavior, we conducted a second-level analysis including the Susceptibility Index (SI) as a covariate. Using the duration-corrected contrasts allowed us to isolate regions in which residual mismatch-related activity predicted behavioral interference. Notably, correlations were observed exclusively in the left hemisphere. Several clusters showed positive correlations with SI, including the cerebellar vermis, left IFG, SMG, and MFG, indicating that more susceptible individuals exhibit stronger neural responses to the feedback mismatch itself (**Table S4, Figure S3B**). The PreCG also remained significant, though located more dorsally compared to the cluster identified prior to the PMod analysis (**Figure 3A**).”

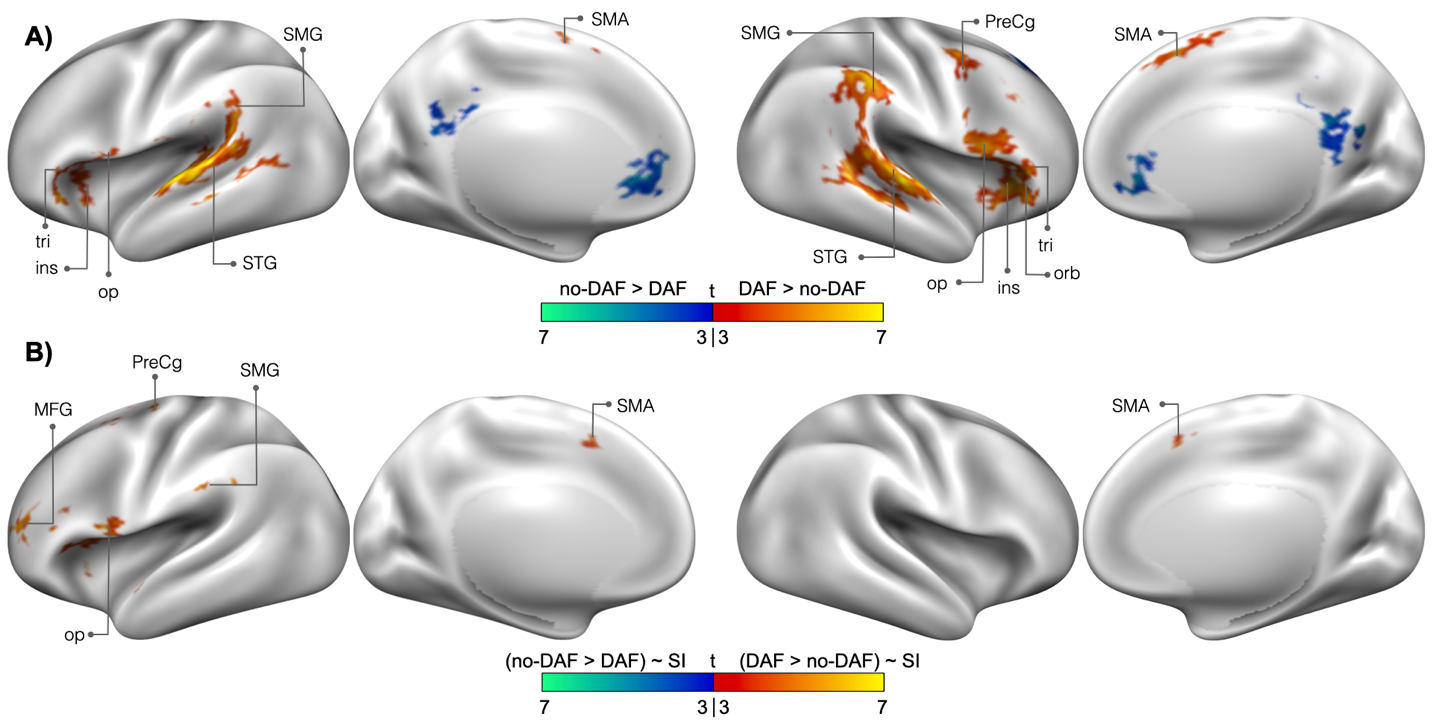

**Supplementary Figure 3:** **Control analysis accounting for speech duration using hierarchical PMods.**

**(A) Main effect of feedback condition (DAF > no-DAF).** Activation maps represent the unique variance associated with the Delay condition after the variance explained by Word Duration was removed via Gram-Schmidt orthogonalization in SPM12. Persistent clusters in the right STG, IFG, and SMG indicate a specialized role in feedback monitoring independent of motor execution length.

**(B) Correlation between duration-corrected activation and Susceptibility Index (DAF > no-DAF ~ SI).** This represents a second-level covariate analysis where the duration-corrected contrast images (DAF > no-DAF) were regressed against each participant's SI. Significant positive correlations were observed primarily in the left hemisphere (IFG, SMG, PreCG, and MFG) and bilaterally in the SMA, suggesting that higher behavioral susceptibility is more closely related to increased neural sensitivity to feedback error than to differences in motoric workload.

Significant clusters identified in SPM12 (p < 0.001 voxel-level; p < 0.05 cluster-level FWE-corrected).

SMA, supplementary motor area; PreCG, precentral gyrus; STG, superior temporal gyrus; SMG, supramarginal gyrus; IFG op/tri/orb, inferior frontal gyrus pars opercularis/triangularis/orbitalis; ins, insula.

**Supplementary Table 3:** Significant fMRI clusters (voxel-level p < 0.001; cluster-level p < 0.05, FWE-corrected) showing greater activation for delayed auditory feedback (DAF > no-DAF) and for immediate feedback (no-DAF > DAF) during speech production, after correcting for speaking duration.

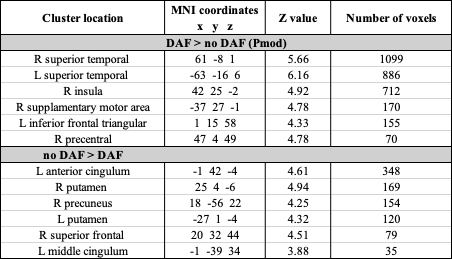

**Supplementary Table 4:** Significant fMRI clusters (voxel-level p < 0.001; cluster-level p < 0.05, FWE-corrected) showing correlations between neural activation (DAF > no-DAF and no-DAF > DAF) and individual susceptibility to DAF (SI), after correcting for speaking duration.

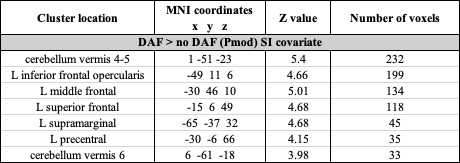

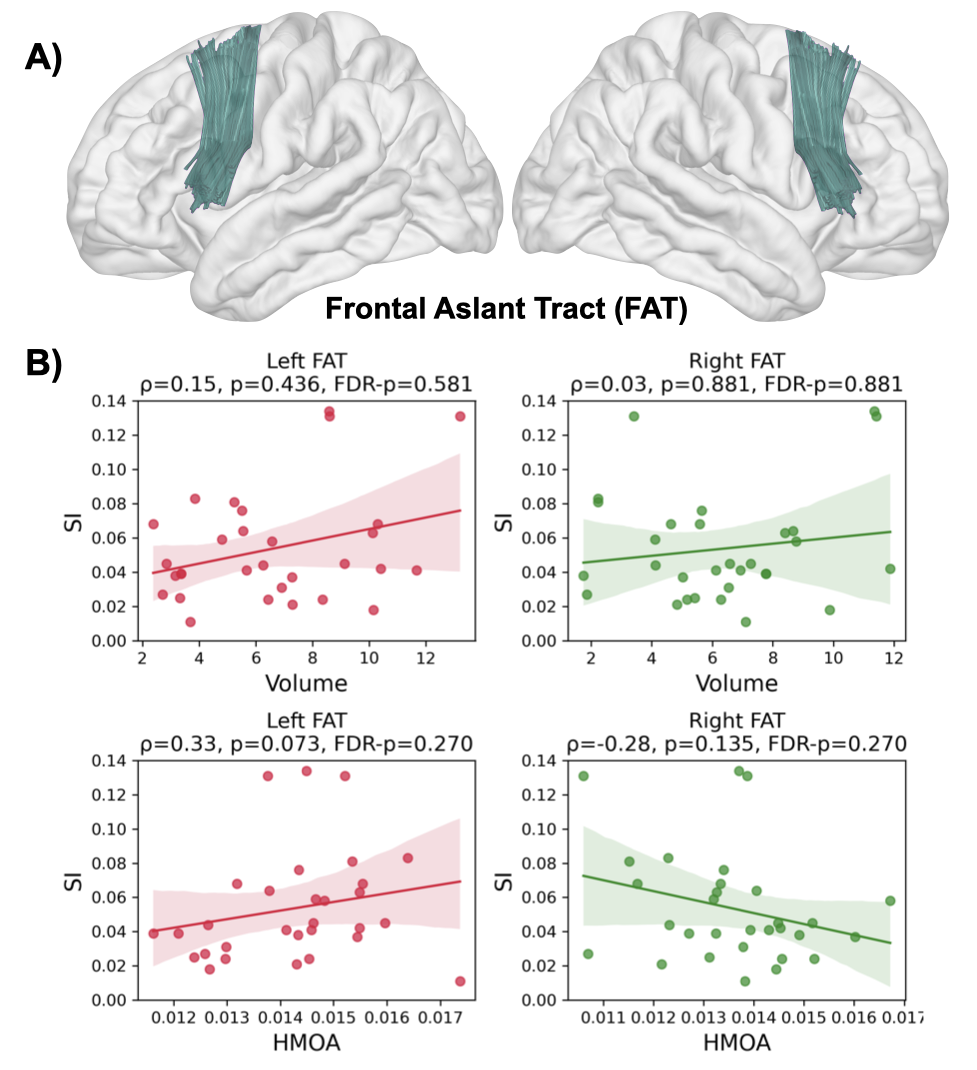

***Supplementary Figure 4: Correlations between individual susceptibility to DAF (SI) and the structural integrity of the frontal aslant tract (FAT).***

***(A)*** *Tractography-based dissection of the frontal aslant tract (FAT)in left and right hemispheres* ***(B)*** *Associations (Spearman correlations) between SI and FAT volume and HMOA are shown for the right (red scatter plots) and left (green scatter plots) hemispheres; points are participants, lines show fitted trend with 95% CI. No significant correlations were observed. Spearman rho, p-value and FDR corrected p-values are shown on top of the plots.*

*HMOA, hindrance-modulated orientational anisotropy; SI, Susceptibility index.*

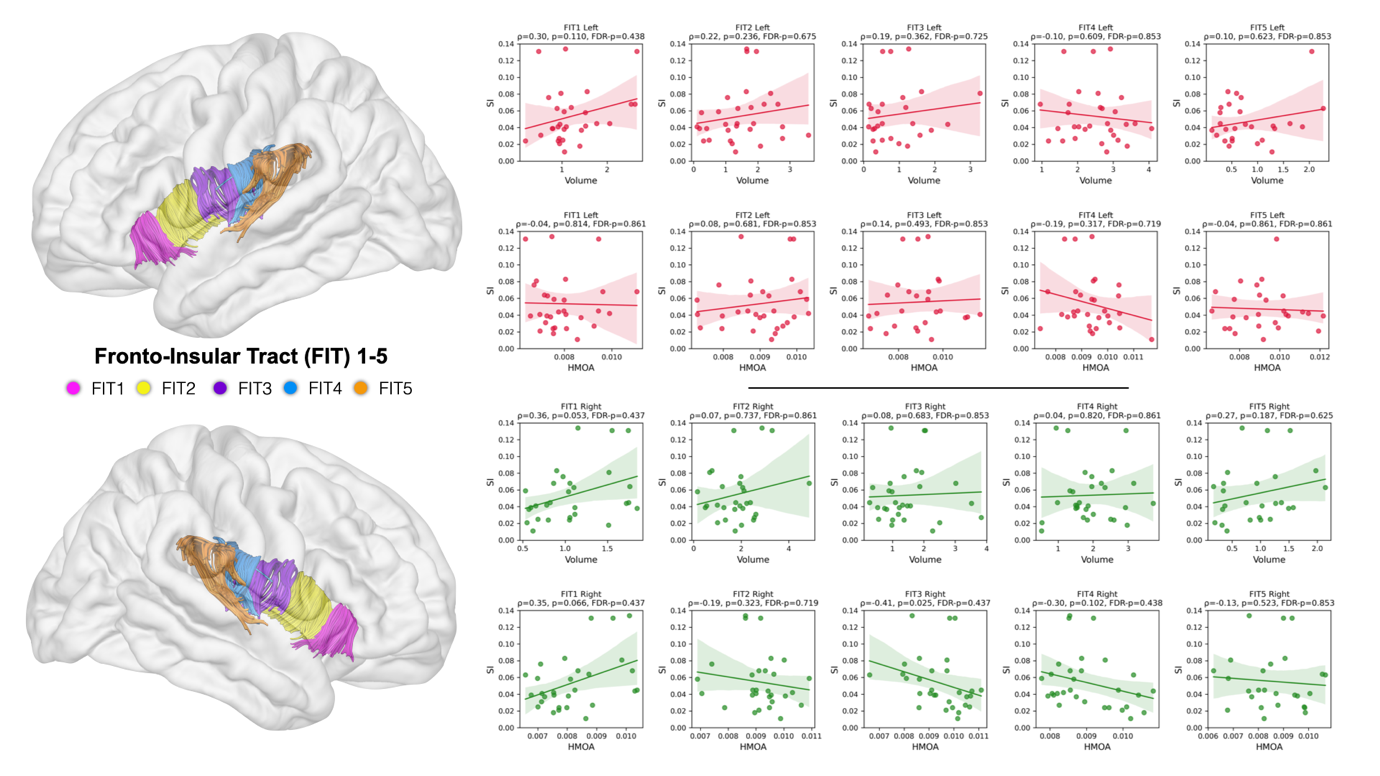

***Supplementary Figure 5: Correlations between individual susceptibility to DAF (SI) and the structural integrity of the five frontal insular tracts (FIT).***

*Tractography-based dissection of the five frontal insular tracts (FIT): FIT1 (anterior insula-pars orbitalis), FIT2 (anterior insula-pars triangularis), FIT3 (anterior insula-pars opercularis), FIT5 (anterior insula-precentral gyrus) and FIT5 (posterior insula-subcentral gyrus). Associations (Spearman correlations) between SI and FIT volume and HMOA are shown for the right (red scatter plots) and left (green scatter plots) hemispheres; points are participants, lines show fitted trend with 95% CI. No significant correlations were observed. Spearman rho, p-value and FDR corrected p-values are shown on top of the plots.*

*HMOA, hindrance-modulated orientational anisotropy; SI, Susceptibility index.*

### ***Supplementary Figure 6: Auditory cortex activation during speech production and passive listening.***

### *fMRI activation is presented for three conditions: speech production with immediate feedback (no DAF; purple), speech production with delayed feedback (DAF; teal), and passive listening to pre-recorded speech (charcoal). Violin plots illustrate the distribution of beta estimates (in arbitrary units) for each condition. Values are calculated relative to an implicit baseline (unmodeled null trials) for six primary and secondary auditory Regions of Interest (ROIs) extracted from the Harvard Oxford atlas (Desikan et al., 2006): Heschl’s Gyrus (HG), Planum Temporale (PT), and posterior Superior Temporal Gyrus (pSTG) bilaterally. Individual subject data points are represented by black dots (strip plot), and black bars indicate the mean with 95% confidence intervals.*

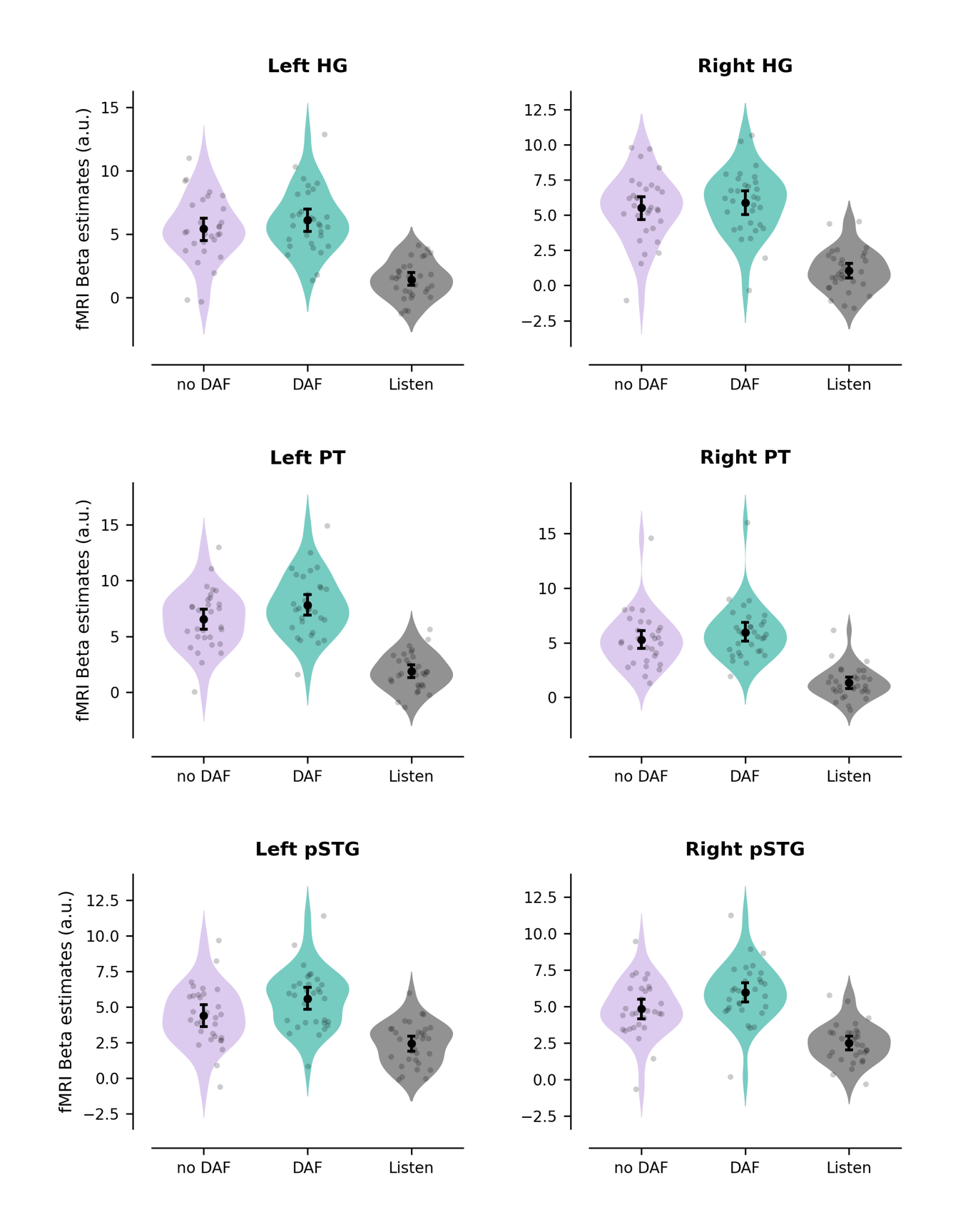

**fMRIPrep BOILERPLATE:**

Results included in this manuscript come from preprocessing performed using *fMRIPrep* 23.1.0 (Esteban et al. (2019); Esteban et al. (2018); RRID:SCR_016216), which is based on *Nipype* 1.8.6 (K. Gorgolewski et al. (2011); K. J. Gorgolewski et al. (2018); RRID:SCR_002502).

**Anatomical data preprocessing**

A total of 1 T1-weighted (T1w) images were found within the input BIDS dataset. The T1-weighted (T1w) image was corrected for intensity non-uniformity (INU) with N4BiasFieldCorrection (Tustison et al. 2010), distributed with ANTs (version unknown) (Avants et al. 2008, RRID:SCR_004757), and used as T1w-reference throughout the workflow. The T1w-reference was then skull-stripped with a *Nipype* implementation of the antsBrainExtraction.sh workflow (from ANTs), using OASIS30ANTs as target template. Brain tissue segmentation of cerebrospinal fluid (CSF), white-matter (WM) and gray-matter (GM) was performed on the brain-extracted T1w using fast (FSL (version unknown), RRID:SCR_002823, Zhang, Brady, and Smith 2001). Brain surfaces were reconstructed using recon-all (FreeSurfer 7.3.2, RRID:SCR_001847, Dale, Fischl, and Sereno 1999), and the brain mask estimated previously was refined with a custom variation of the method to reconcile ANTs-derived and FreeSurfer-derived segmentations of the cortical gray-matter of Mindboggle (RRID:SCR_002438, Klein et al. 2017). Volume-based spatial normalization to one standard space (MNI152NLin2009cAsym) was performed through nonlinear registration with antsRegistration (ANTs (version unknown)), using brain-extracted versions of both T1w reference and the T1w template. The following template was selected for spatial normalization and accessed with *TemplateFlow* (23.0.0, Ciric et al. 2022): *ICBM 152 Nonlinear Asymmetrical template version 2009c* [Fonov et al. (2009), RRID:SCR_008796; TemplateFlow ID: MNI152NLin2009cAsym].

**Functional data preprocessing**

For each of the BOLD runs found per subject (across all tasks and sessions), the following preprocessing was performed. First, a reference volume and its skull-stripped version were generated by aligning and averaging 1 single-band references (SBRefs). Head-motion parameters with respect to the BOLD reference (transformation matrices, and six corresponding rotation and translation parameters) are estimated before any spatiotemporal filtering using mcflirt (FSL , Jenkinson et al. 2002). BOLD runs were slice-time corrected to 0.452s (0.5 of slice acquisition range 0s-0.905s) using 3dTshift from AFNI (Cox and Hyde 1997, RRID:SCR_005927). The BOLD time-series (including slice-timing correction when applied) were resampled onto their original, native space by applying the transforms to correct for head-motion. These resampled BOLD time-series will be referred to as *preprocessed BOLD in original space*, or just *preprocessed BOLD*. The BOLD reference was then co-registered to the T1w reference using bbregister (FreeSurfer) which implements boundary-based registration (Greve and Fischl 2009). Co-registration was configured with six degrees of freedom. First, a reference volume and its skull-stripped version were generated using a custom methodology of *fMRIPrep*. Several confounding time-series were calculated based on the *preprocessed BOLD*: framewise displacement (FD), DVARS and three region-wise global signals. FD was computed using two formulations following Power (absolute sum of relative motions, Power et al. (2014)) and Jenkinson (relative root mean square displacement between affines, Jenkinson et al. (2002)). FD and DVARS are calculated for each functional run, both using their implementations in *Nipype* (following the definitions by Power et al. 2014). The three global signals are extracted within the CSF, the WM, and the whole-brain masks. Additionally, a set of physiological regressors were extracted to allow for component-based noise correction (*CompCor*, Behzadi et al. 2007). Principal components are estimated after high-pass filtering the *preprocessed BOLD* time-series (using a discrete cosine filter with 128s cut-off) for the two *CompCor* variants: temporal (tCompCor) and anatomical (aCompCor). tCompCor components are then calculated from the top 2% variable voxels within the brain mask. For aCompCor, three probabilistic masks (CSF, WM and combined CSF+WM) are generated in anatomical space. The implementation differs from that of Behzadi et al. in that instead of eroding the masks by 2 pixels on BOLD space, a mask of pixels that likely contain a volume fraction of GM is subtracted from the aCompCor masks. This mask is obtained by dilating a GM mask extracted from the FreeSurfer’s *aseg* segmentation, and it ensures components are not extracted from voxels containing a minimal fraction of GM. Finally, these masks are resampled into BOLD space and binarized by thresholding at 0.99 (as in the original implementation). Components are also calculated separately within the WM and CSF masks. For each CompCor decomposition, the *k* components with the largest singular values are retained, such that the retained components’ time series are sufficient to explain 50 percent of variance across the nuisance mask (CSF, WM, combined, or temporal). The remaining components are dropped from consideration. The head-motion estimates calculated in the correction step were also placed within the corresponding confounds file. The confound time series derived from head motion estimates and global signals were expanded with the inclusion of temporal derivatives and quadratic terms for each (Satterthwaite et al. 2013). Frames that exceeded a threshold of 0.5 mm FD or 1.5 standardized DVARS were annotated as motion outliers. Additional nuisance timeseries are calculated by means of principal components analysis of the signal found within a thin band (*crown*) of voxels around the edge of the brain, as proposed by (Patriat, Reynolds, and Birn 2017). The BOLD time-series were resampled into standard space, generating a *preprocessed BOLD run in MNI152NLin2009cAsym space*. First, a reference volume and its skull-stripped version were generated using a custom methodology of *fMRIPrep*. All resamplings can be performed with *a single interpolation step* by composing all the pertinent transformations (i.e. head-motion transform matrices, susceptibility distortion correction when available, and co-registrations to anatomical and output spaces). Gridded (volumetric) resamplings were performed using antsApplyTransforms (ANTs), configured with Lanczos interpolation to minimize the smoothing effects of other kernels (Lanczos 1964). Non-gridded (surface) resamplings were performed using mri_vol2surf (FreeSurfer).

Many internal operations of *fMRIPrep* use *Nilearn* 0.10.1 (Abraham et al. 2014, RRID:SCR_001362), mostly within the functional processing workflow. For more details of the pipeline, see [the section corresponding to workflows in *fMRIPrep*’s documentation](https://fmriprep.readthedocs.io/en/latest/workflows.html).

Copyright Waiver

The above boilerplate text was automatically generated by fMRIPrep with the express intention that users should copy and paste this text into their manuscripts *unchanged*. It is released under the [CC0](https://creativecommons.org/publicdomain/zero/1.0/) license.
